# Female-specific and dosage selections restore genes through transpositions onto the degenerated songbird W chromosomes

**DOI:** 10.1101/692194

**Authors:** Luohao Xu, Qi Zhou

## Abstract

Sex chromosomes are usually suppressed for homologous recombination, which leads to the loss of functional genes on the Y or W chromosomes. It remains unclear how species like birds with a ZW sex system cope with the consequential gene expression imbalance, usually in the absence of global dosage compensation mechanism. Here we tackle this conundrum by reporting 14 genes recently transposed from the Z to the W chromosomes of three songbird lineages, after analyzing a total of 12 songbird species’ genomes. These transpositions are estimated to have occurred within 9 million years. Besides the expected signatures of functional degeneration in some genes on the non-recombining W chromosomes, the other retained genes after transposition are enriched for haploinsufficient genes or housekeeping genes. Several genes show biased expression in ovaries of birds or lizard, or function in female germ cells. These results, together with the reported X-to-Y transpositions provide direct evidence that sex-specific and dosage selections may have recurrently driven the restoration of genes on the Y or W chromosomes, and suggest their evolutionary processes are more dynamic than simply becoming completely degenerated.

---

Y chromosome embarks on an irreversible trajectory of functional degeneration, at regions where its homologous recombination with the X chromosome was suppressed. This is demonstrated by the widely-observed difference of genomic and epigenomic compositions between X and Y chromosomes: while the X is euchromatic and gene rich, the Y chromosome usually has lost most of its functional genes and become highly heterochromatic (Bachtrog 2013; Hughes, et al. 2015). Similar patterns have been found on the W chromosomes in species like birds and butterflies, which have a pair of ZZ chromosomes in males, and ZW chromosomes in females. The divergent evolutionary trajectories between sex chromosome pair are proposed to be driven by the selection for restricting the sex-determining (SD) genes or genes beneficial to one sex but detrimental to the other (so-called ‘sexual antagonistic’, SA genes) within one sex from being inherited in the opposite sex through recombination (Charlesworth and Charlesworth 2000; Ponnikas, et al. 2018). The consequential cost of maintaining sex is essentially a much compromised level of natural selection on the Y/W chromosome due to the lack of recombination(Charlesworth and Charlesworth 2000). This creates a conundrum that when recombination was initially suppressed, the affected regions must contain many other genes with important functions besides the SD/SA genes.

A direct resolution to such ‘collateral damage’ is evolution of dosage compensation on the X/Z chromosome, so that the balance of expression level can be restored. On the other hand, studies showed that the Y/W chromosomes also come up with various strategies to ‘rescue’ functions of certain genes during their complex and dynamic evolutionary course. Some genes with important regulatory functions or high dosage-sensitivity have been demonstrated to be degenerating much slower than others on the mammalian Y (Bellott, et al. 2014; Cortez, et al. 2014) or avian W chromosomes (Smeds, et al. 2015; Bellott, et al. 2017; Xu, et al. 2019) due to a much higher level of selective constraints. The human Y chromosome has evolved palindromic sequence structures to repair deleterious mutations and facilitate gene conversions between Y-linked genes (Rozen, et al. 2003). Other ways of rescuing or even innovating the gene functions on the Y chromosomes include escaping onto the autosomes through transposition(Hughes, et al. 2015), or recruiting novel genes onto the Y chromosomes from various resources. Emerging cases of such gene movements on the Y chromosome have been reported since the characterization of ‘X-transposed, XTR’ region on the male-specific region of human Y chromosome (MSY) over 30 years ago (Page, et al. 1984; Schwartz, et al. 1998; Skaletsky, et al. 2003). This region was transposed from the X chromosome within 4.7 million years (MY)(Ross, et al. 2005) after the human-chimpanzee split, and subsequently disrupted into two blocks by a Y-linked inversion (Schwartz, et al. 1998). The enclosed *PCDH11* X-Y gene pair has been suggested to contribute to the human-specific cerebral asymmetry and language development (Crow 2002; Speevak and Farrell 2011). More cases of transposition from X chromosome or autosomes to the Y chromosome have been reported in Drosophila (Koerich, et al. 2008; Carvalho, et al. 2015; Tobler, et al. 2017) or other Diptera species (Mahajan and Bachtrog 2017), dog (Li, et al. 2013), cat (Li, et al. 2013; Brashear, et al. 2018) and horse (Janečka, et al. 2018), suggesting such transposition events are not rare during the Y chromosome evolution.

Little is known about how the avian W chromosomes resolve the conundrum of losing dosage sensitive genes, particularly without global dosage compensation ever evolved on the homologous Z chromosome (Itoh, et al. 2007; Graves 2014; Gu and Walters 2017). Little genomic information is available except for the euchromatic parts of W chromosomes of chicken (Bellott, et al. 2017), and tens of other bird species (Zhou, et al. 2014; Smeds, et al. 2015; Xu, et al. 2019), although a previous study suggested palindrome structure also exists on the W chromosomes of sparrows and blackbirds (Davis, et al. 2010). One might expect transposition or retrotransposition events are scarce in avian genomes due to their compact structures with a much lower repeat content to mediate these events, particularly the L1 retroposons relative to mammals (International Chicken Genome Sequencing 2004; Suh 2015). Indeed, there are only 51 retrogenes identified in chicken, relative to over 15,000 cases in mammals(International Chicken Genome Sequencing 2004). So far no transposed genes have been reported on the avian W chromosomes, and we have recently reported one retrotransposed gene on a songbird W chromosome (Xu, et al. 2019). Of course, these results are far from being conclusive regarding the role of transposition or retrotransposition in the evolution of avian W chromosomes, because only a few out of over 10,000 bird species have been investigated. In addition, the degree of sexual selection, which is known to dramatically vary across bird species, must have a different impact shaping the evolution of sex chromosomes.

Here we looked into this question by studying 12 songbird genomes which both male and female sequencing data is available. We reasoned that these Illumina-based genomes do not contain complete information of complex sequence structures (e.g., palindromes) or traces of ancient transposition events on the W chromosome. We therefore focused on identifying the recent transpositions, if any from the Z onto the W chromosome that are manifested as female-specific elevations of both read coverage and heterozygosity level. While other regions with such an elevation pattern in both sexes are inferred as Z-linked duplications, those at the end of the chromosome with elevation of female coverage to the rest Z-linked regions, but without sex-biased patterns of heterozygosity are inferred as pseudoautosomal regions (PAR) (Figure 1, Supplementary Figure 1). Intriguingly, we identified four Z-to-W transposition events involving 14 genes among great tit (*Parus major*), medium ground finch (*Geospiza fortis*), red bird-of-paradise (*Paradisaea rubra*) and Raggiana bird-of-paradise (*P. raggiana*). The two birds-of-paradise species share the same transposition, and for simplicity hereafter we used red bird-of-paradise to represent this lineage. The lengths of detected transposed regions range from 67kb in great tit to 1.3Mb in red bird-of-paradise. We dated the transposition of medium ground finch about 8.3 million years (MY) ago, as the same transpositions have been found in all the other Coerebinae (Darwin’s finches and their relatives) but absent in their sister group Sporophilinae (Lamichhaney, et al. 2015) (Supplementary Figure 2). Similarly, we dated the transpositions of red bird-of-paradise within 4MY (Supplementary Figure 3) and that of great tit about 7 MY ago, after examining their sister species.

**Fig. 1.**
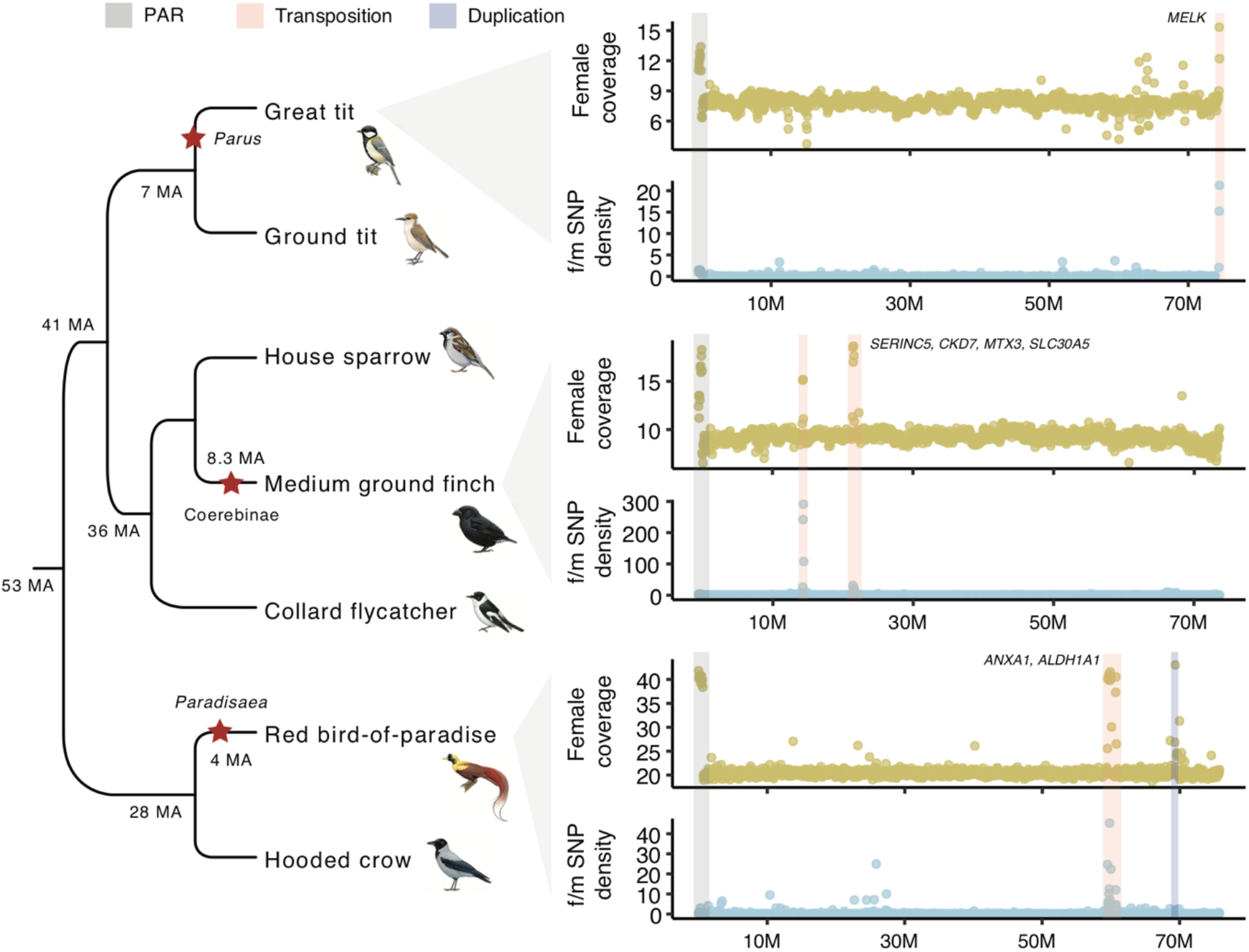
Transpositions from the Z to W chromosomes in songbirds. Genomic regions on the Z chromosome showing female-specific elevations of SNP density and coverage were inferred as recent transposition event. Pseudoautosomal regions (PARs) and Z-linked duplications do not show elevated SNP density. We show seven representative species out of the 12 studied songbirds, including three species with signatures of transpositions. The red asterisks indicate the origination branch of the transpositions.

These very recent Z-to-W transpositions provide a unique window for us to examine the evolution of W-linked genes at their early stages. They show clear signatures of functional degeneration. For instance, among the five genes transposed in medium ground finch, at least one (*THBS4*) has become a probable pseudogene due to frameshift mutations. The most prominent case of gene loss after transposition is found in red bird-of-paradise. Almost half of the originally transposed sequences, involving a large 583kb region and 4 encompassing genes and 2 partial genes, and a nearby smaller 2kb region (Supplementary Figure 4) have become deleted, with the deleted regions showing similar levels of coverage and heterozygosity with other non-transposed Z-linked regions in the female (Figure 2, Supplementary Note). Based on analyses of insert size of mate-pair libraries, we have not identified any large-scale insertions into the transposed regions.

**Fig. 2.**
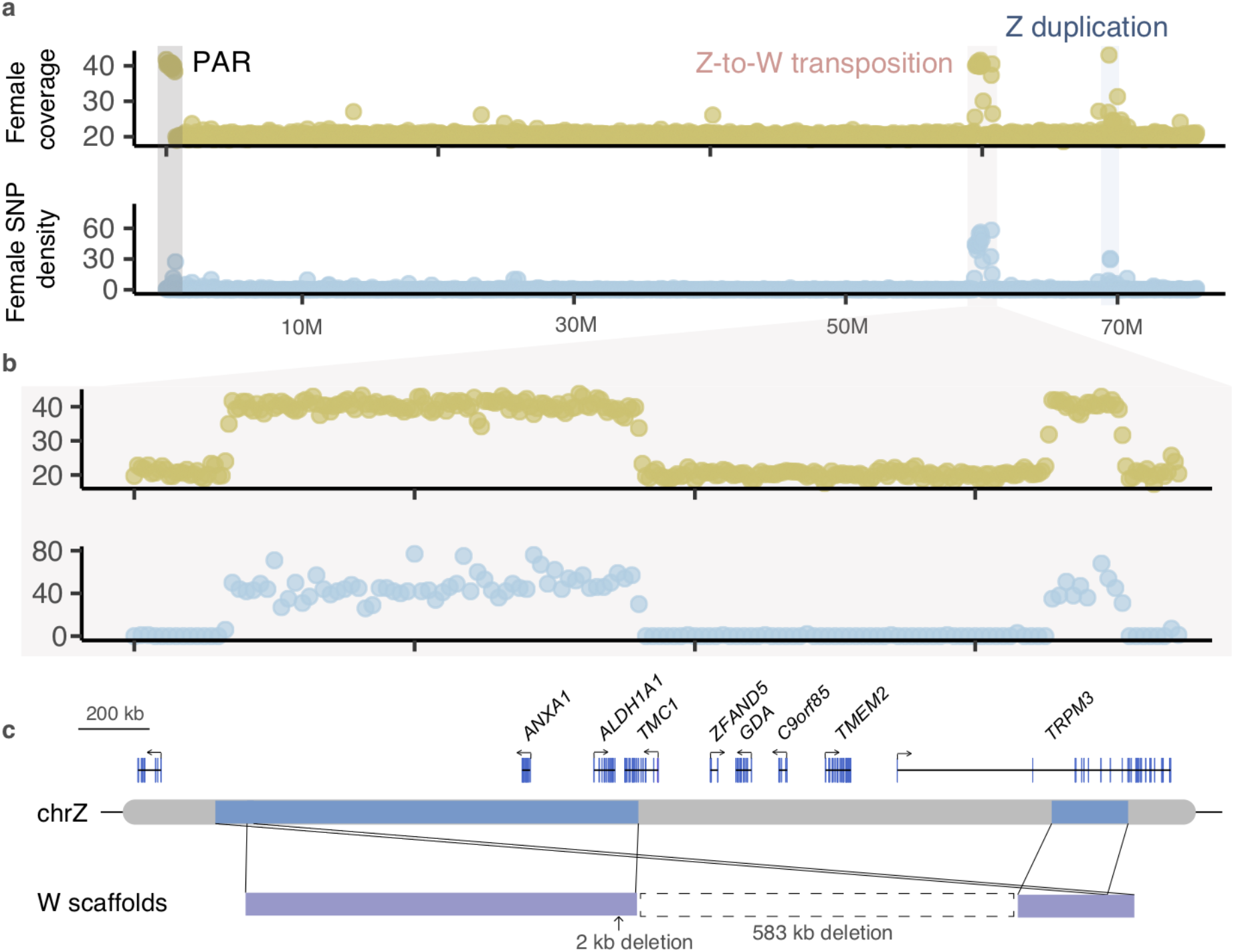
The Z-to-W transposition in red bird-of-paradise. a) the loci of transposition (at ~60 Mb) shows an elevated heterozygosity and coverage in females. b)-c) a zoom-in view of the transposed region. The 1.3 Mb transposed sequence contains 9 genes, but 5 compete and 2 partial genes probably have become lost through a 583 kb sequence deletion. Only *ANXA1* and *ALDH1A1* are retained on the W.

While such gene losses are expected because of the lack of recombination, the retained genes, essentially the recently restored genes that had previously become lost on the W chromosomes, are more informative for the driving forces that originally fixed these transpositions. We reasoned that two types of selection, i.e., female-specific selection for the female reproductive genes, as well as dosage selection for the haploinsufficient genes probably account for the restoration of W-linked genes. The first type of selection is demonstrated by a previous study showing that the chicken breeds selected for higher female fecundity exhibit an increased W-linked gene expression than other breeds (Moghadam, et al. 2012). Indeed, the only two retained genes *ANXA1* and *ALDH1A1* after the transposition in red bird-of-paradise (Figure 2), and the great tit transposed gene *MELK* all have a biased or specific expression pattern in ovary in many examined bird species (Supplementary Figure 5), and also their outgroup species green anole lizard (Figure 3). Although *ALDH1A1* has a relatively lower expression in ovary than in testis, it has been recently shown in mice that the disruption of this gene delays the onset of meiosis in ovary (Bowles, et al. 2016). Besides, *ANXA1* and *CDK7* probably have been restored by strong dosage selection, indicated by their much higher levels of predicted haploinsufficiency (HP score) than most other genes on the Z chromosome (Supplementary Figure 5) (Huang, et al. 2010), as well as a lack of any nonsynonymous changes compared to their Z-linked homologs (Supplementary Table 1). Several medium ground finch genes, for example, *SERINC5* and *MTX3*, have a low HP score, but a very broad expression pattern across tissues measured by tissue-specificity matrix *tau*, thus are likely restored as housekeeping genes (Figure 3). In fact, the restored genes have a generally higher HP score (*P*=0.051, Wilcoxon test) than the genes that have become lost after the transpositions.

**Fig. 3.**
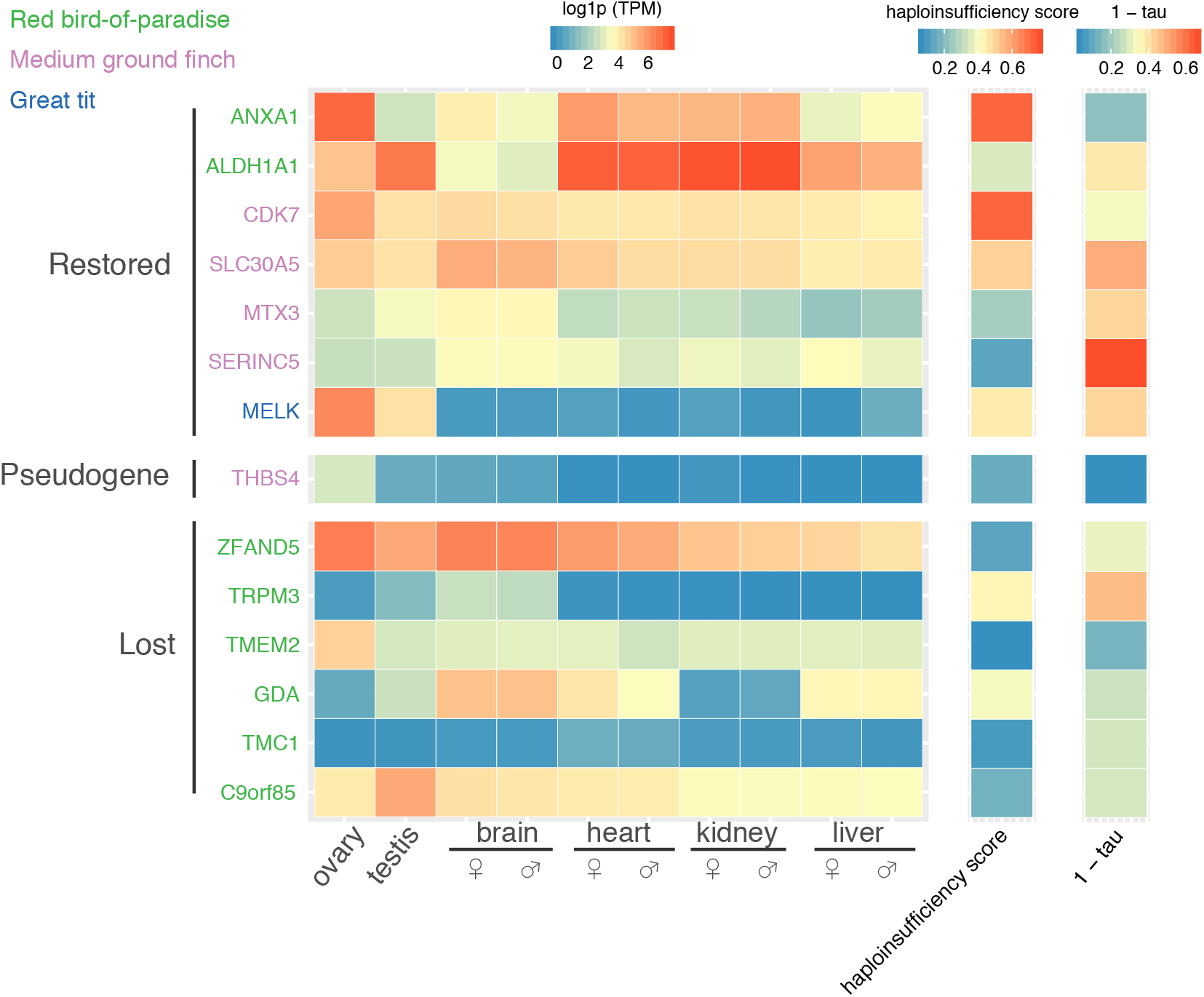
Female-specific and dosage selections restore avian W-linked genes. The seven restored genes through transposition on the W chromosomes tend to show a higher expression level or a broader (larger 1-tau value) expression pattern across tissues than the lost genes. Most of restored genes also have a higher degree of dosage sensitivity (higher haploinsufficiency scores) than the lost genes, with some genes (e.g., *ANXA1*) showing an ovary-biased expression pattern.

These results together provide clear evidence for the female-specific and dosage selections have driven the frequent restoration of W-linked genes through transpositions among songbird species. Because similar X-to-Y transpositions have been reported in insects and mammals, we propose that restoration of once-lost genes onto the non-recombining sex chromosomes is probably a general feature in sex chromosomes evolution. Although such restoration is not expected to alter the evolutionary trajectories of W or Y chromosomes toward complete functional degeneration, in fact, we found some transposed genes have already become lost or show signatures of functional degeneration (e.g., *THBS4*). Such loss-and-restoration cycles may recurrently occur throughout the evolution of sex chromosomes, particularly in ZW systems that usually do not have global dosage compensation to cope with the imbalance of gene expression. We have to point out that our method can only identify recent transpositions, and probably has missed ancient transpositions that have become too divergent in sequence between Z and W chromosomes. The genes involved in the such cases nevertheless have probably already become pseudogenes. Our results are in line with the reported cases in avian W or mammalian Y chromosomes that dosage-sensitive genes are retarded for their functional degeneration due to the strong selective constraints (Bellott, et al. 2014; Smeds, et al. 2015; Bellott, et al. 2017; Xu, et al. 2019). We also provided new evidence that sex-specific selection is shaping the evolution of the W chromosome, which was assumed to be less frequent than that shaping the Y chromosome, due to the more frequent and intensive male-targeted sexual selection.

## Materials and Methods

The genomic, transcriptomic and resequencing data used in this study are listed in Supplementary Table 2-4. For the 12 songbird genomes, genomic data are available for both sexes except for three species. We first used the published Z chromosome sequence of great tit (Laine, et al. 2016) to identify and order the Z-linked sequences among the investigated species (Supplementary Figure 6, Supplementary Note). To calculate the read coverage, we first mapped the reads to the reference genomes using BWA-MEM (0.7.16a-r1181) with default parameters. We used the function ‘depth’ in samtools (1.9) to calculate coverage for every nucleotide site, subsequently removed those sites with mapping quality (-Q) lower than 60 or depth 3 times higher than average. Then we calculated genomic coverage of every 50 kb sliding window by using ‘bedtools map’ function. Any windows with less than 60% of the region (30 kb) mapped by reads were excluded. We used the GATK (3.8.0) pipeline (HaplotypeCaller) to call variants. Raw variants were filtered by this criteria: -window 10 -cluster 2 “FS > 10.0”, “QD < 2.0”, “MQ < 50.0”, “SOR > 1.5”, “MQRankSum < −1.5”, “ReadPosamplenkSum < −8.0”. We further required the variants showing an allele frequency between 0.3 and 07 (the expected heterozygosity should be 0.5 for one individual). The SNP density was defined by the number of SNPs over a 50 kb window. To genotype the W-derived alleles, we used the FastaAlternateReferenceMaker to create W-linked sequences for the transposed regions. The gene models on the W were then predicted by genewise (2.4.1). To remove potential chimeric W-derived alleles in the Z-linked regions (due to the collapse of genome assembly), if any, we used male sequencing reads to polish the Z-linked sequence using pilon (1.22). To estimate pairwise substitution rated between sex-linked alleles, we used the guidance program (v2.02) and PRANK (170427) to align the Z- and W-linked coding sequences. Then we used the ‘free ratio’ model in codeml from PAML package (4.9e) to estimate the substitution rates. We used the program RSEM (1.3.0) to estimate gene expression levels. Details of the method is described in Xu et al. (2019). Codes used in this study has been deposited at Github (https://github.com/lurebgi/ZWtransposition)

## Acknowledgment

We would like to thank Fumin Lei and Yalin Chen at the Institute of Zoology, Chinese Academy of Sciences for sharing their unpublished genomic data of the green-backed tit (*Parus monticolus*) for inferring the origination time of transposition in great tit. L.X. is supported by the uni:docs fellowship programme from University of Vienna. Q.Z. is supported by the National Natural Science Foundation of China (31722050, 31671319), the Fundamental Research Funds for the Central Universities (2018XZZX002-04), and start-up funds from Zhejiang University.

